# Description of *Gloeomargarita ahousahtiae* sp. nov., a thermophilic member of the order Gloeomargaritales with intracellular carbonate inclusions

**DOI:** 10.1101/2022.11.03.515036

**Authors:** Thomas Bacchetta, Purificación López-García, Ana Gutiérrez-Preciado, Neha Mehta, Feriel Skouri-Panet, Karim Benzerara, Maria Ciobanu, Naoji Yubuki, Rosaluz Tavera, David Moreira

## Abstract

A unicellular cyanobacterium, strain VI4D9, was isolated from thermophilic microbial mats thriving in a hot spring of the Ahousaht territory of Vancouver Island, Canada. The cells were elongated rods (5.1 μm in length and 1.2 μm in width on average). Their UV-visible absorption spectra revealed that they contain chlorophyll *a*, phycocyanin, and carotenoids. Transmission electron microscopy showed the presence of thylakoids concentrated on one side of the cells. The strain grew within a temperature range of 37–50°C, with an optimum at 45°C. Its genome had a size of 3,049,282 bp and a DNA G+C content of 51.8 mol%. The cells contained numerous intracellular spherical granules easily visible under scanning electron microscopy. Energy-dispersive x-ray spectroscopy revealed that these granules were made of Ca-, Ba- and Sr-containing carbonates. A phylogenetic 16S rRNA gene tree robustly placed this strain as sister to several environmental sequences and the described species *Gloeomargarita lithophora*, also characterized by the possession of intracellular carbonate inclusions. We consider strain VI4D9 to represent a new *Gloeomargarita* species based on its marked phenotypic differences with *G. lithophora*, notably, its thermophilic nature and different thylakoid organization. We propose the name *Gloeomargarita ahousahtiae* sp. nov. for this newly isolated thermophilic cyanobacterium. The type strain is VI4D9 (Culture Collection of Algae and Protozoa strain 1472/1; Laboratorio de Algas Continentales Mexico strain LAC 140). *G. ahousahtiae* is the second species described within the recently discovered order Gloeomargaritales.

## Introduction

The Great Oxygenation Event that took place about 2.3 billion years ago (Bekker & Holland, 2004) entailed a considerable change in Earth’s ecosystems. Cyanobacteria were responsible for atmospheric oxygenation, being the only lineage performing oxygenic photosynthesis at that time (Soo *et al*., 2017). Nowadays, oxygen production is mainly shared between cyanobacteria and photosynthetic eukaryotes, including green plants and a large diversity of algae. The first photosynthetic eukaryotes to evolve, the Archaeplastida (including red algae, glaucophytes, and land plants + green algae) emerged through a symbiosis that involved a cyanobacterial endosymbiont and a heterotrophic eukaryotic host (Moreira & Philippe, 2001; Keeling, 2013). This event, called primary plastid endosymbiosis, was at the origin of plastids and allowed eukaryotes to perform oxygenic photosynthesis via their cyanobacterial-derived plastids. This capability was later transferred to other eukaryotic lineages through secondary and tertiary endosymbioses involving green and red algae as endosymbionts (Keeling, 2013; McFadden, 2001; Ponce-Toledo *et al*., 2019). The monophyly of Archaeplastida has been found in phylogenetic trees reconstructed both with plastid- and nucleus-encoded markers (Moreira *et al*., 2000; Rodriguez-Ezpeleta *et al*., 2005; Isarri *et al*., 2021), hence supporting the hypothesis that a single primary endosymbiotic event gave rise to this group. However, the identity and lifestyle of the cyanobacterial endosymbiont involved in this major evolutionary event have long been debated.

Different studies have alternatively proposed early- and late-branching cyanobacteria (e.g., Blank, 2013; Dagan *et al*., 2013) but only recently the cyanobacterial sister group of plastids has been confidently identified. This group is represented by the deep-branching cyanobacterium *Gloeomargarita lithophora* strain Alchichica-D10 (Ponce-Toledo *et al*., 2017), which was isolated from microbialite samples from Lake Alchichica in Mexico (Couradeau *et al*., 2012; Moreira *et al*., 2018). Phylogenetic analysis of conserved protein markers supported that *G. lithophora* defined a new cyanobacterial order, the Gloeomargaritales (Moreira *et al*., 2018). *G. lithophora* exhibits the unusual ability to synthesize large amounts of intracellular amorphous calcium carbonate inclusions, sometimes enriched in barium and strontium (Couradeau *et al*., 2012; Benzerara *et al*., 2014). While cyanobacteria have been known for long to induce extracellular calcium carbonate precipitation as a consequence of the environmental pH increase triggered by photosynthesis (Riding, 2006), this intracellular biomineralization process was only recently discovered in several cyanobacterial lineages (Cam *et al*., 2018; Benzerara *et al*., 2022).

Moreover, *G. lithophora* has a unique preference to incorporate Ba, followed by Sr and lastly Ca within the intracellular carbonate inclusions (Cam *et al*., 2016; Mehta *et al*., 2022). Up to now, *G. lithophora* was the only isolated species within the Gloeomargaritales. Nevertheless, environmental surveys identified a large diversity of related 16S rDNA sequences in diverse freshwater environments, mostly microbialites and thermophilic microbial mats (Ragon *et al*., 2014). Moreover, *Gloeomargarita*-like cells containing intracellular carbonates have also been observed by electron microscopy in microbial mat samples of the Meskoutine hot spring in Algeria (Amarouche-Yala *et al*., 2014), although they have never been grown in the laboratory. Here, we describe the second isolated Gloeomargaritales strain: *Gloeomargarita ahousahtiae* strain VI4D9.

## Materials and methods

### Sampling site and isolation

Thermophilic microbial mat samples were collected in sterile plastic containers in August 2005 in Hot Springs Cove (Vancouver Island, Canada). In their natural habitat, these mats are irrigated by a continuous flow of sulfur-rich, moderately hot (45-47 °C) and circumneutral (pH 7.5) water. Since that sampling date, the mats were maintained alive in laboratory aquaria at 45 °C under a 12h-12h light-dark cycle.

To isolate new non-filamentous cyanobacterial species from the biofilms growing in these aquaria, we first resuspended the biofilm cells by vortexing and repeated pipetting, and subsequently filtered the cell suspension through a 5 μm pore size filter. Filtrate volumes ranging from 0.5 to 2 μl were then used to inoculate three 96-well microplates containing BG-11 medium. Microplates were incubated at 45 °C for several months under a dark-light (12h-12h) cycle. Wells with cyanobacterial growth (identified by their blue-green coloration) were serially diluted in BG-11 until pure cultures were obtained.

### DNA extraction, 16S rRNA gene amplification and genome sequencing

Cyanobacterial cells were collected by centrifugation of liquid cultures at 10000 g for 5 min and DNA was extracted from cell pellets with the DNeasy PowerBiofilm kit (Qiagen) following manufacturer’s instructions.

16S rRNA genes were amplified by PCR using the two Gloeomargaritales-specific primers 69F-Gloeo (AAGTCGAACGGGGKWGCAA) and 1227R-Gloeo (GATCTGAACTGAGACCAAC) (Ragon *et al*., 2014). PCR reactions were done in 25 μl of reaction buffer, containing 1 μl of the eluted DNA, 1.5 mM MgCl2, dNTPs (10 nmol each), 20 pmol of each primer, and 0.2 U Taq platinum DNA polymerase (Invitrogen). PCR reactions were run under the following conditions: 35 cycles (denaturation at 94 °C for 15 s, annealing at 55 °C for 30 s, extension at 72 °C for 2 min) preceded by a 2-min denaturation step at 94 °C, and followed by a 7-min extension step at 72 °C. Positive amplicons were sequenced using the same amplification primers (Beckman Coulter Genomics, Takeley, UK).

The 16S rRNA gene sequence was used to identify a new cyanobacterial species from the order Gloeomargaritales (see results). Its DNA was extracted from a liquid culture as explained above and sequenced using both short (paired-end Illumina HiSeq 2500) and long (Nanopore MinION) reads. A hybrid assembly of short and long reads was produced with Unicycler v0.4.9b (Wick *et al*., 2017). The assembled genome was annotated with Prokka v1.14.5 (Seeman, 2014). To compare different genomes, we used average nucleotide identity (ANI), estimated using the EZBioCloud ANI Calculator (Yoon *et al*., 2017), and the digital DNA-DNA hybridization (dDDH), estimated using the GGDC calculator (Meier-Kolthoff *et al*., 2021).

### Optical and electron microscopy

Phase contrast and differential interference contrast (DIC) microscopy observations were done using a Zeiss Axioplan 2 Imaging light microscope (Jena, Thuringia, Germany). Pictures were taken with both an AxiocamMR camera using the Zeiss AxioVision 4.8.2 SP1 suite and a Sony α9 (Minato, Tokyo, Japan) digital camera.

We applied two different scanning electron microscope (SEM) techniques. On the one hand, we observed dehydrated cells previously filtered on 0.22 μm polyethersulfone (PES) filters, rinsed with MilliQ water and dried at ambient temperature. These cell-covered PES filters were mounted on aluminum stubs using double-sided carbon tape and carbon-coated prior to SEM observation with a Zeiss Ultra55 SEM microscope (Jena, Thuringia, Germany). On the other hand, we observed fresh non-dehydrated cells deposited on 0.22 μm PES filters using a Hitachi SU5000 FEG Low Vacuum microscope (Tokyo, Japan).

The chemical composition of intracellular carbonate inclusions was studied using scanning transmission electron microscopy (STEM) and an energy dispersive x-ray spectrometer (EDXS). STEM analyses were performed in the high-angle annular dark-field (HAADF) mode using a JEOL 2100F microscope (Akishima, Tokyo, Japan) operating at 200 kV and equipped with a field emission gun and a JEOL EDXS detector. For STEM observations, cyanobacterial cells were harvested by centrifugation at 10000 g (5 min) and the cell pellets were washed twice before resuspension in 500 μl of Milli-Q water and deposited on pre-ionized carbon-coated 200-mesh copper grids.

We prepared ultrathin sections for transmission electron microscopy (TEM) analysis of cells collected by centrifugation (10000 g, 5 min) and resuspended in a solution of 2.5% glutaraldehyde and 2% paraformaldehyde in 0.2 M sodium cacodylate buffer (pH 7.2). After rinsing the cells with the buffer, the suspension was fixed in 1% osmium tetroxide, followed by dehydration through an ethanol bath series of 10 min each at concentrations of 30%, 50%, 70%, 90%, followed by 3 baths of 10 min in 100% ethanol before final substitution with acetone. The fixed cells were embedded in low viscosity resin (Agar Scientific) and thin sections were prepared with a diamond knife mounted on a Leica UC6 ultramicrotome (Wetzlar Germany) and observed with a JEOL JEM 1400 microscope (Akishima, Tokyo, Japan).

### Pigment characterization

We collected cells by centrifugation (10000 g, 5 min) and purified pigments from the cell pellet by 100% acetone overnight extraction at 4 °C. The absorption spectrum of different dilutions of the pigments was measured with a Hach DR 5000 spectrophotometer (Ontario, Canada) in the wavelength range from 350 to 800 nm.

## Results

### Isolation of a new Gloeomargaritales cyanobacterium

We collected samples of thermophilic microbial mats growing on a small, moderately hot (45-47 °C) and circumneutral (pH 7.5) stream in the Hot Springs Cove hydrothermal system (Supplementary fig. S1) in Vancouver Island (Canada). Since the collection date (August 2005), the mats were incubated alive in laboratory aquaria at 45 °C and with a 12h-12h light-dark cycle. After several months of incubation, the walls of the aquaria were covered by a conspicuous green biofilm (Supplementary fig. S2).

Previous work had shown that cyanobacterial species belonging to the order Gloeomargaritales can be very diverse in moderately thermophilic microbial mats (Amarouche-Yala *et al*., 2014; Ragon *et al*., 2014). To isolate new Gloeomargaritales species from our aquaria biofilm, we resuspended it by strong vortexing and pipetting and inoculated 96-well microplates containing BG-11 medium with the cell suspension after filtering it through a 5 μm pore size filter. After several months of incubation at 45 °C, we observed two wells with growth of cells with typical cyanobacterial pigmentation. Microscopy observation confirmed the presence of rod-shaped cyanobacterial cells, but also of *Chloroflexus-like* bacteria and some filamentous cyanobacteria. PCR amplification and sequencing of 16S rRNA genes from DNA extracted from these wells using Gloeomargaritales-specific primers showed the presence of a new Gloeomargaritales species. It was closely related to several 16 rRNA gene environmental sequences obtained from various hot springs of the Yellowstone National Park (Fig. 1). This group of sequences from hydrothermal systems branched as sister to the group containing the only Gloeomargaritales species described so far, *Gloeomargarita lithophora*. We used serial dilution to obtain a pure culture (strain VI4D9), which we named *Gloeomargarita ahousahtiae* (see the Taxonomic analysis below).

**Fig. 1.**
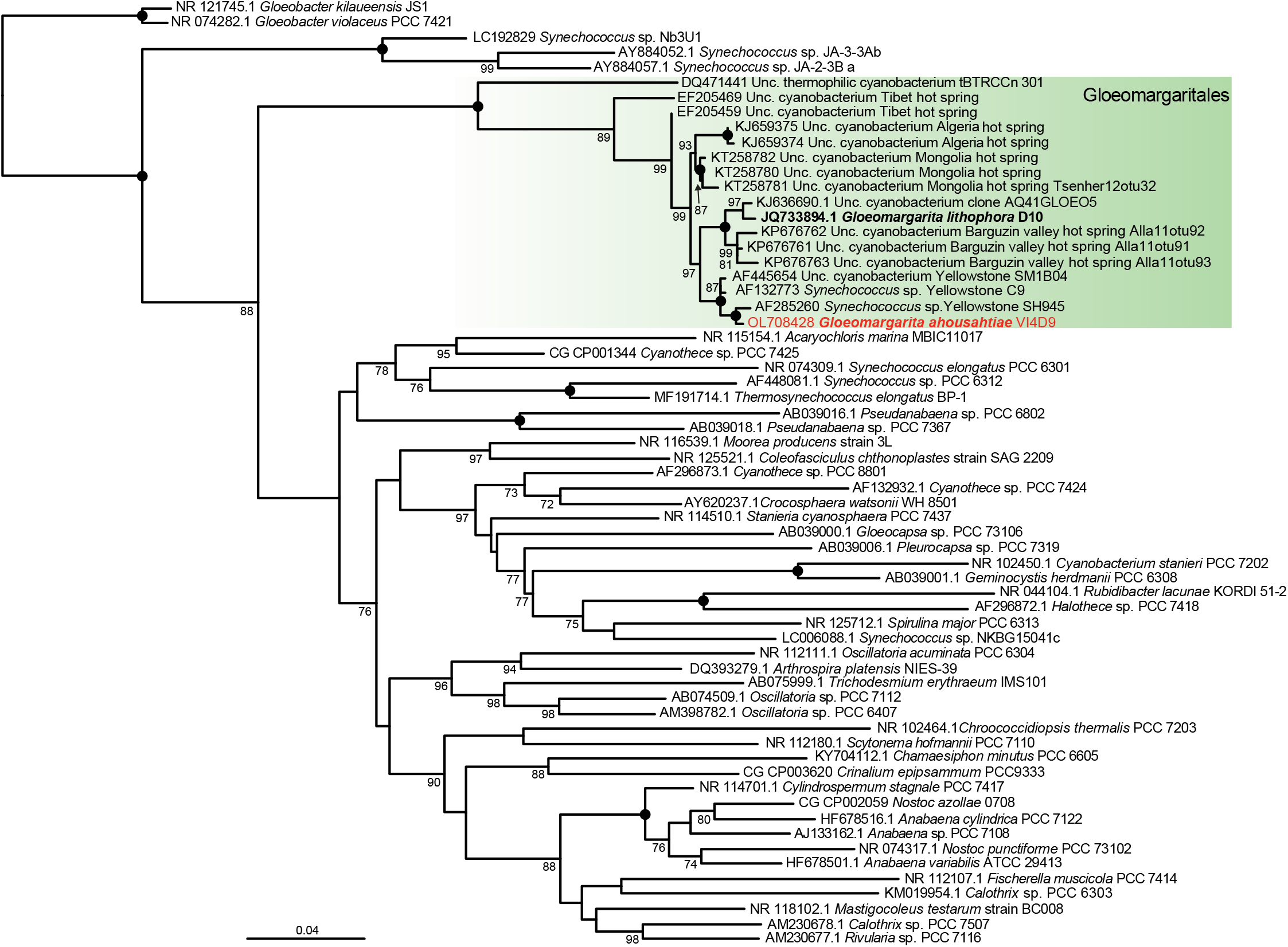
16S rRNA gene maximum likelihood phylogenetic tree of cyanobacteria showing the position of *Gloeomargarita ahousahtiae* within the Gloeomargaritales (green clade). Numbers on branches are bootstrap proportions (only values >70% are shown); black circles indicate 100% bootstrap values.

### Genome characteristics

Using a combination of short (Illumina HiSeq, 30 million reads for a total length of 9.1 Gb) and long (Nanopore, 1.3 million reads for a total length of 20.6 Gb) reads, we were able to assemble the complete genome sequence of *G. ahousahtiae* in a single contig of 3,162,419 bp. The genome had a G+C content of 51.8 mol% and encoded 3,141 genes, including 3,059 protein-coding genes. A single rRNA locus was present, as well as genes for all common tRNAs. 1495 (48.9%) of the protein-coding genes were annotated in comparison with proteins with known biological functions and 1564 (51.1%) remained annotated as hypothetical. These general values were similar to those of the *G. lithophora* genome, which has a size of 3,049,282 bp, a G+C content of 52.2 mol%, and encodes 3,101 genes (Moreira *et al*., 2018). 2463 genes of *G. ahousahtiae* had homologs in the genome of *G. lithophora*. We compared both Gloeomargaritales genomes using the average nucleotide identity (ANI) and the digital DNA-DNA hybridization (dDDH). We obtained values of 82.0% for the ANI and 24.90% for the dDDH.

In addition to complete gene sets coding for the proteins involved in oxygenic photosynthesis and carbon fixation typical of cyanobacteria, the genome of *G. ahousahtiae* contains, among other important features, genes coding for a large set of nitrogenase subunits (including *nifB, nifD, nifE, nifH, nifK, nifN, nifU, nifV, nifW, nifX*, and *nifZ*) as well as a number of ABC transporters for metals and inorganic ions, such as bicarbonate, sulfate, nitrate, molybdate, iron, phosphate, manganese, and cobalt.

### *Phenotypic characteristics of* Gloeomargarita ahousahtiae

Cells grew mainly attached to surfaces and their growth was faster in glass culture flasks than in plastic flasks. Since this strain had a benthic lifestyle and agitation to obtain cultures in suspension was deleterious at all temperatures, no measurement of growth rate and generation time based on culture optical density was possible. Growth was therefore measured as the time that the strain required to cover the bottom of the culture flasks starting from identical inoculum amounts. We examined the growth of the new strain in BG-11 medium at different temperatures ranging from 20 °C to 60 °C. Growth was detected only at temperatures between 37 °C and 50 °C, with an optimum at 45 °C.

*Gloeomargarita ahousahtiae* cells were short rods that measured 5.1±0.9 μm in length and 1.2±0.02 μm in width (from 62 cells measured) (Figs 2-3) and divided by binary fission. These cells were longer than those of *G. lithophora* (3.9±0.6 μm in length and 1.1±0.1 μm in width) (Moreira *et al*., 2018). The cultures were intensely colored (Fig. 4). The absorption spectrum of acetone-extracted *G. ahousahtiae* pigments showed two absorption peaks at 437 and 662 nm typical of chlorophyll *a*, a peak at 616 nm likely corresponding to phycocyanin, and a typical carotenoid peak at 478 nm. (Figs 5-6). This pigment combination is characteristic for the majority of freshwater cyanobacteria (Chen *et al*., 2021).

**Figs 2-3.**
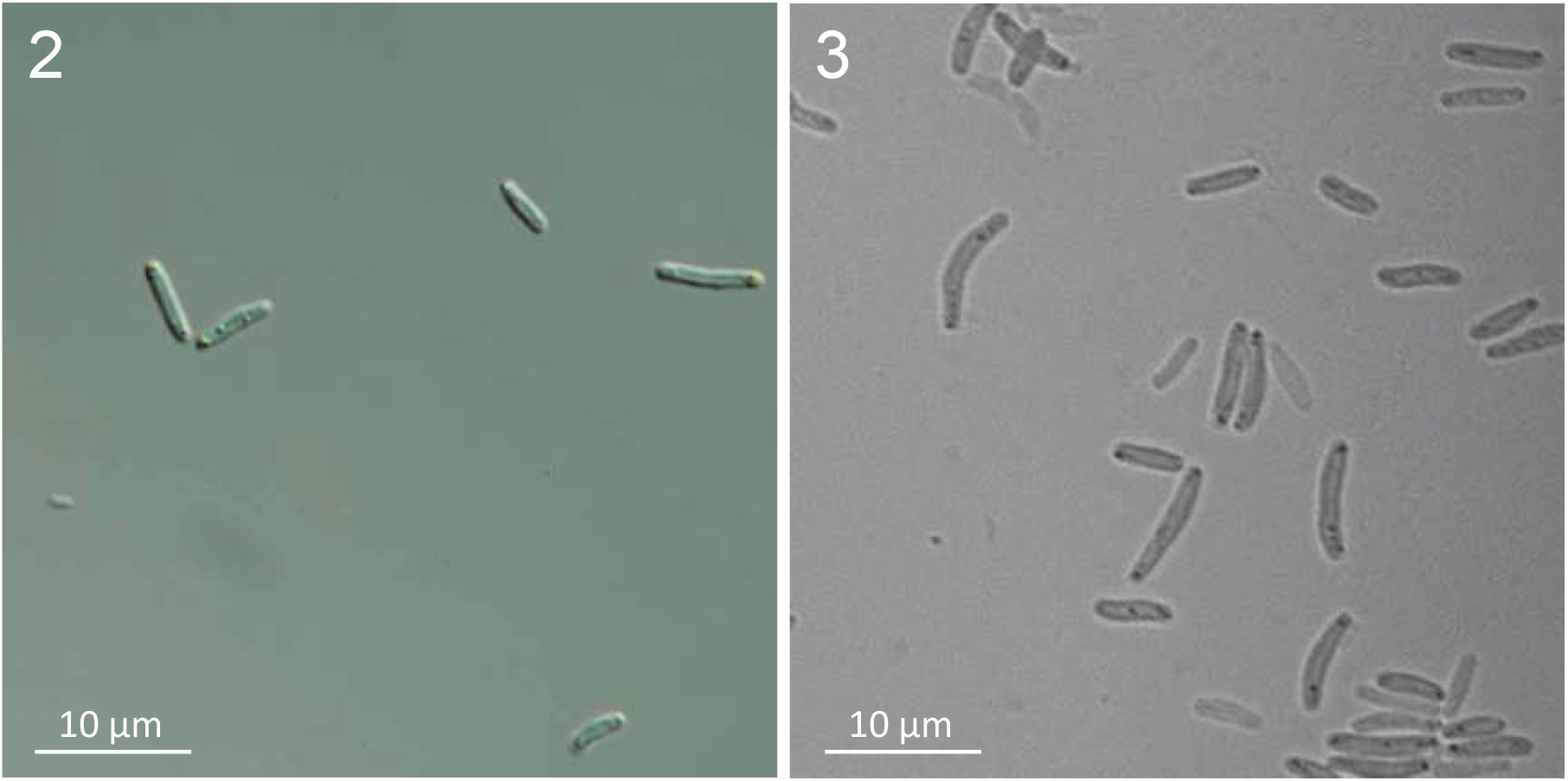
Optical microscopy images of *Gloeomargarita ahousahtiae.* **Fig. 2**, differential interference contrast image. **Fig. 3**, phase contrast image.

**Figs 4-6.**
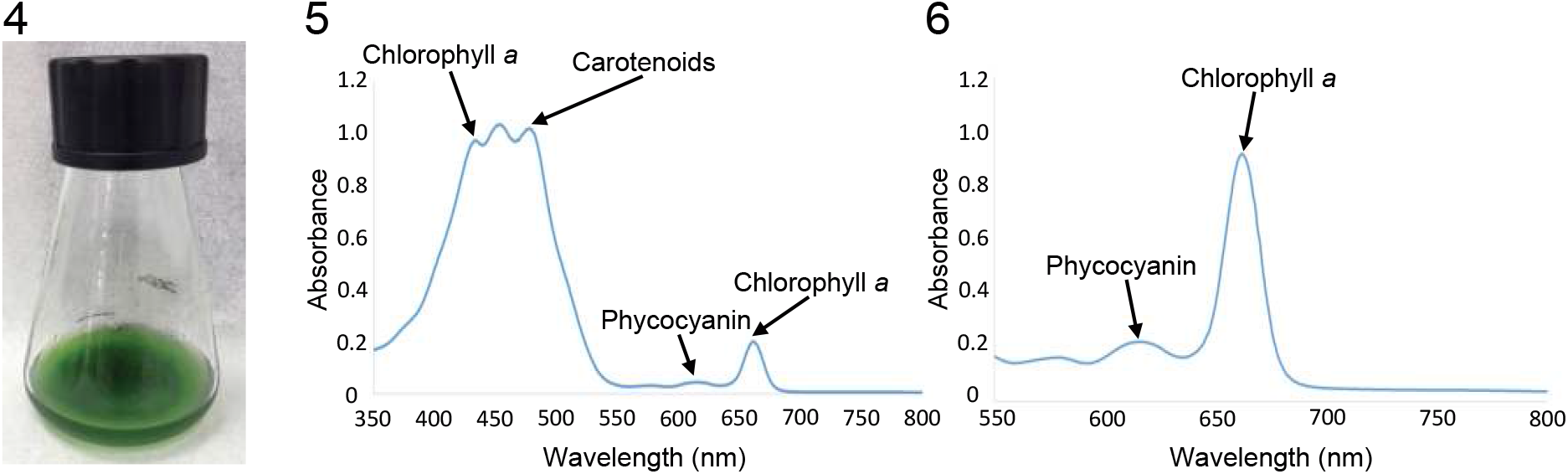
Pigments of *Gloeomargarita ahousahtiae.* **Fig. 4**, view of a liquid culture after one month of growth, showing a typical cyanobacterial color. **Fig. 5**, absorbance spectrum (between 350 and 800 nm) of *G. ahousahtiae* pigments purified by 100% acetone extraction. **Fig. 6**, absorbance spectrum (between 550 and 800 nm) of *G. ahousahtiae* concentrated (4x) pigments to better appreciate the phycocyanin peak.

We observed *G. ahousahtiae* under scanning electron microscopy (SEM) using both dried carbon-coated cells and fresh cells (see Materials and methods). With both approaches we noticed that, similar to *G. lithophora, G. ahousahtiae* forms intracellular amorphous carbonate inclusions scattered within the cytoplasm, albeit often arranged in a rather linear configuration (Figs 7-8). These carbonate inclusions were accompanied by polyphosphate granules, clearly recognizable by their lower brightness under SEM observation (Figs 8-9). Inspection of the cell surface at high magnification showed a finely dotted pattern (Fig. 10).

**Figs 7-10.**
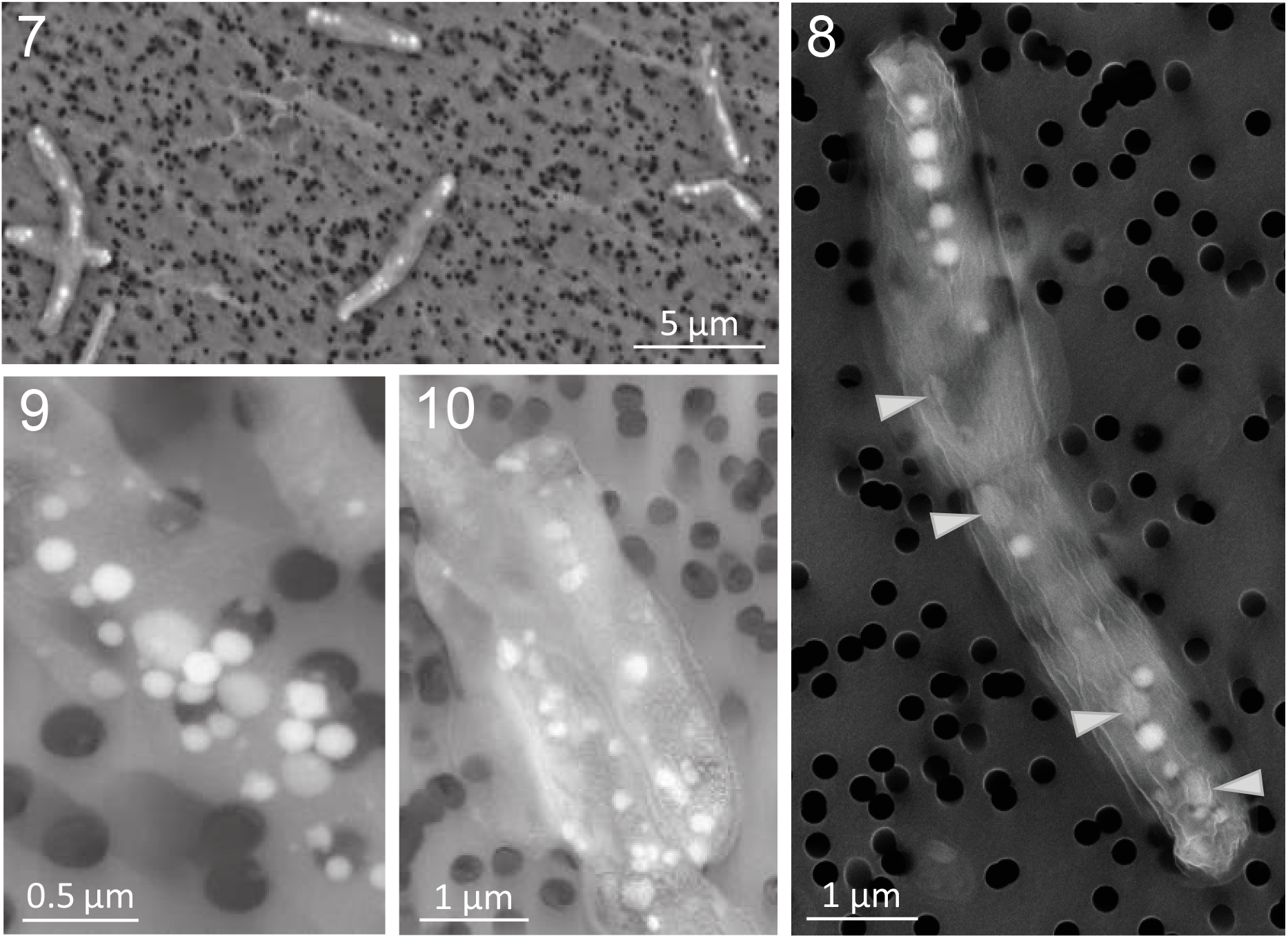
Scanning electron microscopy images of *Gloeomargarita ahousahtiae* cells. **Fig. 7**, general view of several fresh cells observed under low vacuum; notice the bright intracellular carbonate inclusions. **Fig. 8**, image of a dried cell acquired using the secondary electron (SE) mode. In addition to bright carbonate inclusions, the cell also contains a few darker, and generally bigger, polyphosphate granules (white arrowheads). **Fig. 9**, close view of bright (carbonate) and grey (polyphosphate) inclusions in a fresh cell observed under low vacuum with a BSE detector. **Fig. 10**, overlay of images of fresh cells acquired using backscattered (BSE) and secondary (SE) electron detectors under low vacuum; notice the finely dotted cell surface.

To test whether *G. ahousahtiae* also had the ability reported in *G. lithophora* to selectively uptake Ba and Sr into the intracellular carbonate inclusions (Cam *et al*., 2916), we incubated our strain in BG-11 medium amended with Ba and Sr (final concentration 25 μM). After one month of growth, we analyzed the composition of intracellular carbonate inclusions using a scanning transmission electron microscope (STEM) and an energy dispersive x-ray spectrometer (EDXS). The detection of characteristic peaks of Ba, Sr and Ca in the SEM-EDXS spectrum provided the first clue that *G. ahousahtiae* accumulated all these elements within the intracellular inclusions (Fig. 11). Furthermore, STEM-HAADF revealed that some intracellular inclusions had a brighter core surrounded by a darker layer, whereas other inclusions exhibited uniform high brightness (Fig. 12). The EDXS analysis showed that inclusions made only of Ba, Sr or Ca were those that appeared as uniform bright spheres in the STEM-HAADF observations, whereas inclusions that had a Ba or Sr core surrounded by a layer of Sr or Ca were those that showed the differential layered brightness (Figs 13-17). Based on a comparison with what was previously observed for *G. lithophora* (Cam *et al*., 2016), the formation of these layered intracellular carbonates suggested that *G. ahousahtiae* selectively uptakes Ba and Sr over Ca. The cells also contained phosphorus-rich inclusions that were most likely polyphosphates (Fig. 17).

**Figs 11-17.**
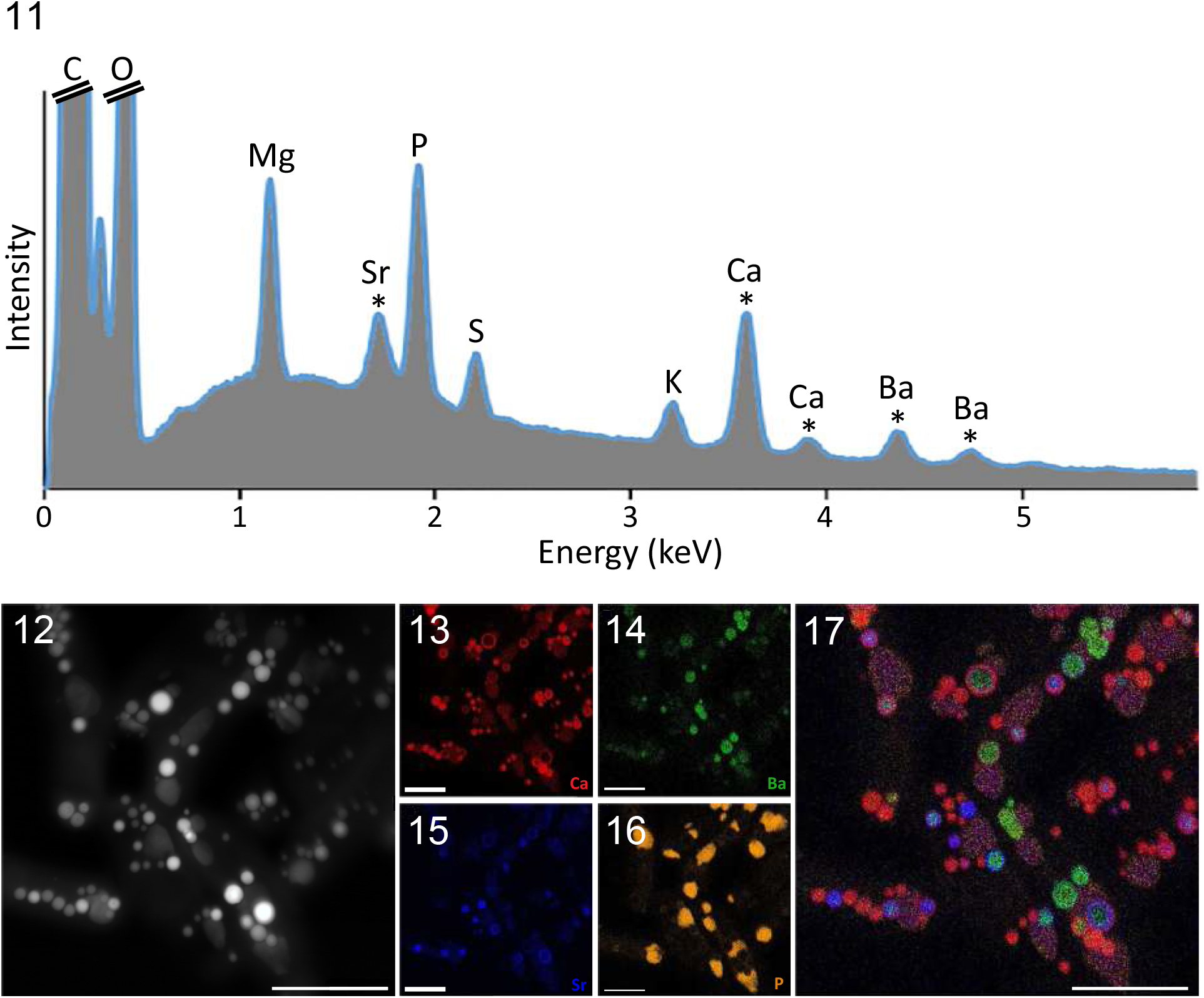
SEM-EDXS and STEM-EDXS analyses of *Gloeomargarita ahousahtiae* cells after one month of growth. **Fig. 11**, EDXS spectrum showing the presence of Ca, Ba and Sr in the cells. **Fig. 12**, STEM-HAADF image of cells showing carbonate (bright white) and polyphosphate (darker) inclusions. **Figs 13-16**, Ca (red), Ba (green), Sr (blue) and P (yellow) EDXS maps of these cells. **Fig. 17**, Overlay of the Ca, Ba, Sr and P EDXS maps. All scale bars correspond to 2 μm.

Transmission electron microscopy (TEM) observation of thin sections showed that *G. ahousahtiae* cells exhibited a typical Gram-negative structure with two membranes and a thin intermediate peptidoglycan wall (Figs 18-19). In contrast with the concentric thylakoids close to the cell membrane found in *G. lithophora* (Moreira *et al*., 2018), those of *G. ahousahtiae* were mainly located along one side of the cell (Fig. 18). Many structures with low electron density were observed in the cytoplasm (Fig. 20), which likely corresponded to empty spaces left by the fast dissolution of the intracellular carbonate inclusions during the sample preparation (Benzerara *et al*., 2014; Li *et al*., 2016).

**Figs 18-20.**
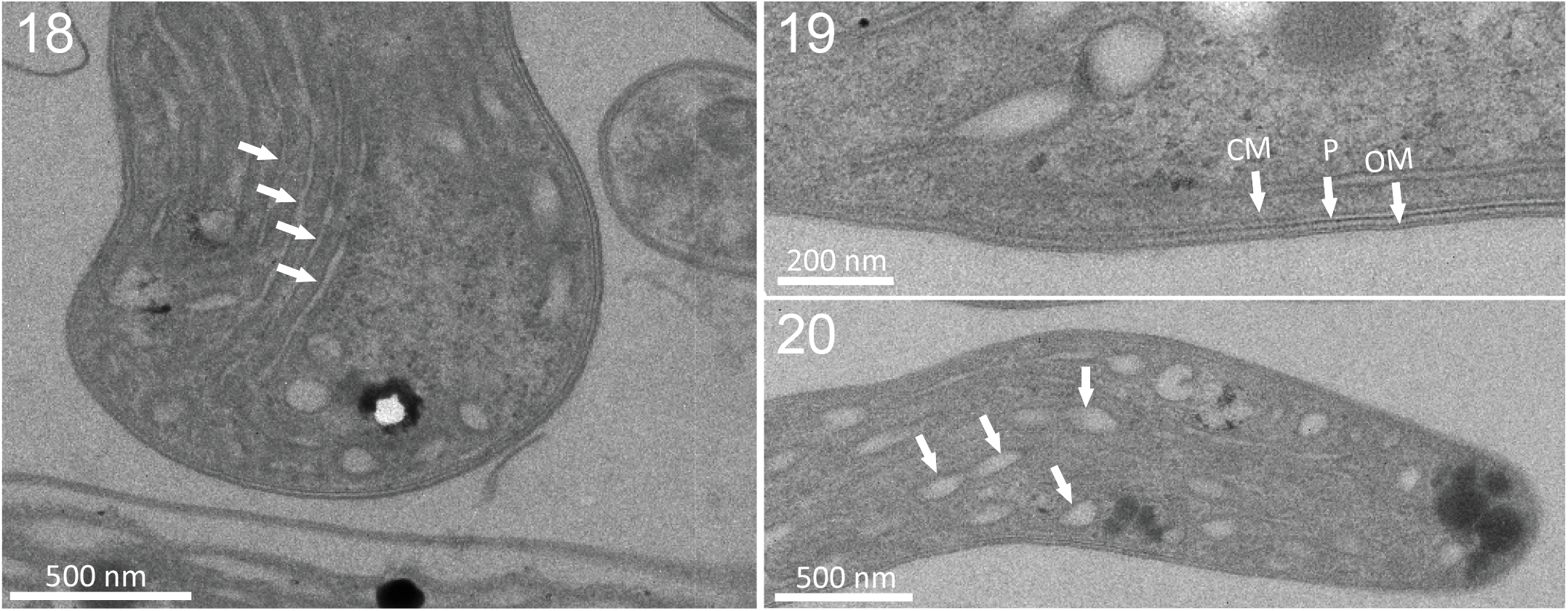
Transmission electron micrographs of thin sections of *Gloeomargarita ahousahtiae* cells. **Fig. 18**, cell section showing thylakoids concentrated on one side of the cell (white arrows). **Fig. 19**, cell section showing the cell (CM) and outer (OM) membranes and the peptidoglycan wall (P) between them. **Fig. 20**, cell section showing numerous intracellular structures with low electron density (white arrows).

### Taxonomic analysis

#### *Gloeomargarita ahousahtiae* T.Bacchetta, P.López-García, & D.Moreira, sp. nov

*Description* Elongated rod-shaped cells with average cell size of 5.1 μm in length and 1.2 μm in width. Shows slow benthic growth on surfaces at temperatures ranging from 37 °C to 50 °C, with an optimum at 45 °C in BG-11 medium. Oxygenic photoautotrophic metabolism. Contains chlorophyll *a*, phycocyanin and carotenoids, and possesses thylakoids mainly located along one side of the cell. Uses Ba, Sr and Ca to form intracellular carbonate spherical inclusions.

HOLOTYPE: (here designated): FIslAho-1, deposited at FCME Herbarium of the Faculty of Sciences at the UNAM, Mexico. Cultured material fixed on microscope slides. Figure 2 illustrates the holotype.

TYPE STRAIN: CCAP 1437/1 and PMC 919.15, deposited at the Culture Collection of Algae and Protozoa (Scottish Association for Marine Science laboratory, UK) and the Laboratorio de Algas Continentales. Ecología y Taxonomía, UNAM, Mexico), respectively.

TYPE LOCALITY: Hot Springs Cove hydrothermal system, Vancouver Island, Canada (49°21’59.99”N, 126°15’27.00”W), microbial mats in a small stream. Sample collected in August 2005.

ETYMOLOGY: The species name refers to the Ahousaht people, the community that occupies and manages the hot springs where the strain comes from.

DNA SEQUENCES: Sequences were deposited in GenBank with accession numbers OL708428 and OV696605.

## Discussion

Despite their phylogenetic proximity (Fig. 1), the two Gloeomargaritales species *G. lithophora* and *G. ahousahtiae* exhibit several important phenotypic differences that reinforce their distinction as separate species. An evident one is the much higher optimal growth temperature of *G. ahousahtiae* (45 °C instead of 30 °C), which makes of this species the first isolated thermophilic representative of the Gloeomargaritales. The 16S rRNA gene sequence of *G. ahousahtiae* is closely related to several environmental sequences from continental hydrothermal systems. In fact, moderate thermophily seems to be the most widespread phenotype among the Gloeomargaritales, as deduced from the diversity of environmental sequences obtained from continental hot springs in various continents (see Fig. 1 and Ragon *et al*., 2014). In that sense, *G. ahousahtiae* constitutes a good representative model to study the biology of this cyanobacterial order.

The two Gloeomargaritales species exhibit other significant differences, for example at the ultrastructural level. Whereas *G. lithophora* has thylakoids arranged as concentric layers beneath the cytoplasmic membrane (Blondeau *et al*., 2018), those of *G. ahousahtiae* appear in most cells concentrated on one side of the cell (Fig. 18). In fact, differences in thylakoid structure are common between mesophilic and thermophilic cyanobacteria, with a tendency to be more irregular in the later (Mares *et al*., 2019). From a metabolic point of view, an interesting difference between both species is the presence of a large set of *nif* genes only in *G. ahousahtiae*, which is comparable to that of nonheterocystous diazotrophic cyanobacteria (e.g., Nonaka *et al*., 2019) and that most likely allows *G. ahousahtiae* to synthetize a functional nitrogenase complex. *G. ahousahtiae* also possesses genes coding for ABC transporters involved in bicarbonate, molybdate, and cobalt uptake, which are absent in *G. lithophora*.

Despite these differences, both *Gloeomargarita* species share a conspicuous similarity, which is the presence of numerous intracellular carbonate inclusions (Figs 7-10). Although this biomineralization process can be found in several other cyanobacterial groups (Benzerara *et al*., 2022), the two *Gloeomargarita* species are unique in their strong preference to use strontium and barium over calcium to synthetize their carbonate inclusions (Cam *et al*., 2016 and this work). Therefore, this seems to be a general trait in the *Gloeomargarita* genus common to both mesophilic and thermophilic species. The putative function of these intracellular carbonate granules remains unknown. One possibility may be that they participate in the control of cell buoyancy by increasing the cell density, which may be especially relevant in benthic species such as the Gloeomargaritales. However, carbonate inclusions are also found, although in lower abundance, in planktonic species such as *Microcystis aeruginosa* (Benzerara *et al*., 2022). Alternatively, these inclusions may be a by-product of photosynthesis, which can increase intracellular pH and induce carbonate precipitation in the presence of bivalent cations such as Ca^2+^. However, this does not explain the marked preference of Gloeomargaritales to incorporate strontium and barium in their carbonates, which suggests the existence of an active mechanism to transport these elements (Cam *et al*., 2016; Benzerara *et al*., 2022). Further research will be necessary to understand the possible role of these intracellular biominerals in cyanobacteria.

## Acknowledgements

We thank Prof. Denis Lynn for help with sampling and the UNICELL single-cell genomics platform (https://www.deemteam.fr/en/unicell) for Nanopore sequencing. This work has benefited from the facilities and expertise of MIMA2 (Université Paris-Saclay, INRAE, AgroParisTech, 78350, Jouy-en-Josas, France.) microscopy platform. We thank Vlad Costache (MIMA2 - Micalis Institute) for help with SEM imagining.

## Author contributions

TB isolated the new strain, carried out molecular, phylogenetic and morphological analyses, and wrote the manuscript draft; MC and AGP contributed to genome sequencing and analysis; NM, FSP and KB took microphotographs and described the chemistry of intracellular carbonate inclusions; NY carried out TEM observations; PLG and DM participated in sampling, supervised and coordinated the work, and wrote the final version of the manuscript. All authors read and approved the final version of the manuscript.

## Supplementary Information

The following supplementary material is accessible via the Supplementary Content tab on the article’s online page at https://doi.org//xxxxxxxxxxxxxxxx

**Supplementary figure S1**. Sampling site at Hot Springs Cove hydrothermal system showing conspicuous thermophilic microbial mats.

**Supplementary figure S2**. Laboratory aquaria incubated at 45°C showing the growth of a cyanobacterial biofilm.

## Data availability

The 16S rRNA gene and complete genome sequences of *Gloeomargarita ahousahtiae* have been submitted to GenBank (accession numbers OL708428 and OV696605, respectively).

