## Supplemental figures for "Description of *Gloeomargarita ahousahtiae* sp. nov., a thermophilic member of the order Gloeomargaritales with intracellular carbonate inclusions"

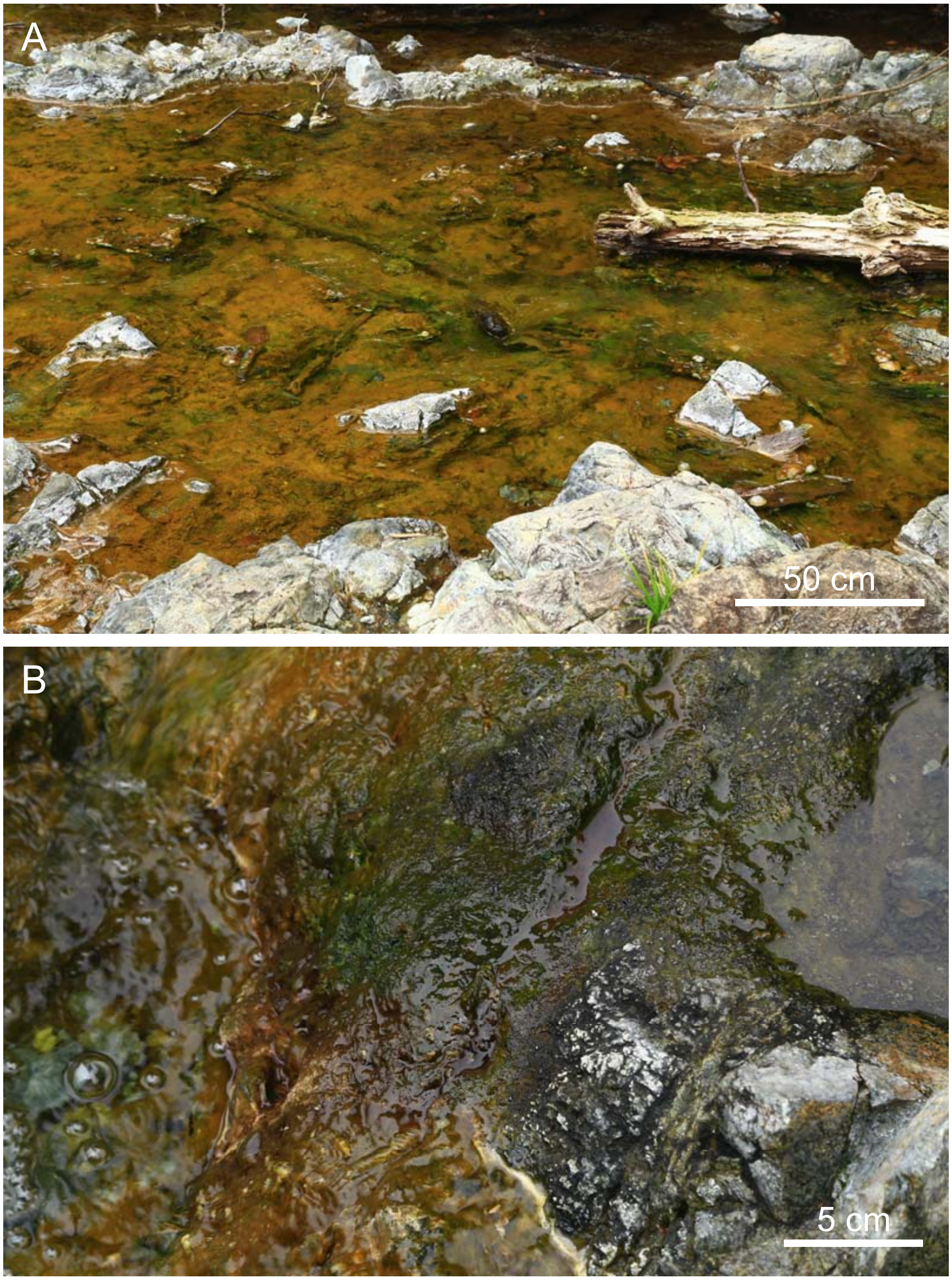


**Supplementary figure S1.** Sampling site. A) General view of microbial mats growing in the the warm stream at Hot Springs Cove. B) Closer view of the microbial mat.

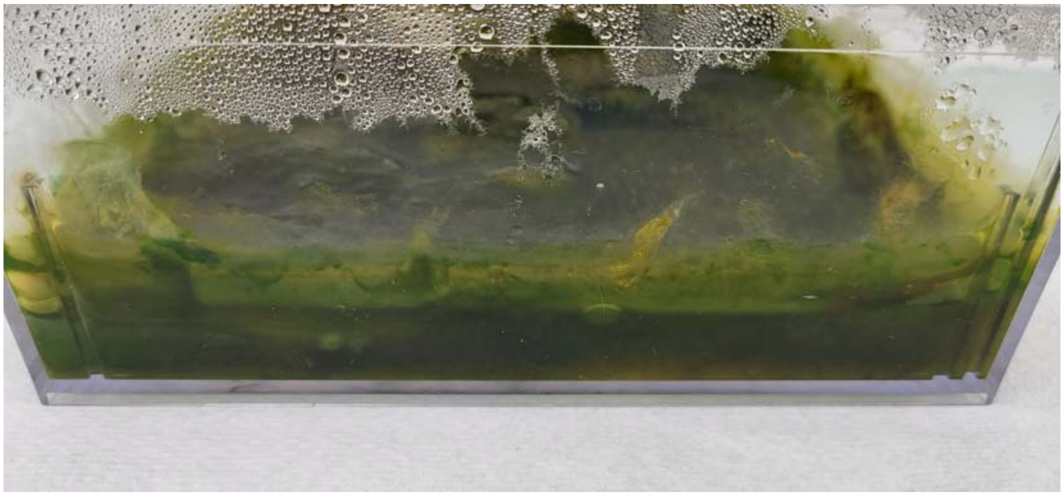

**Supplementary figure S2.** Cyanobacterial biolfilm developping on the wall of an aquarium incubated at 45°C containing mat samples from Hot Springs Cove.
